# Restoring persistent accessibility to memories after sleep deprivation-induced amnesia

**DOI:** 10.1101/2021.05.11.443364

**Authors:** Youri G. Bolsius, Pim R.A. Heckman, Frank Raven, Elroy L. Meijer, Martien J.H. Kas, Steve Ramirez, Peter Meerlo, Robbert Havekes

**Author notes:** Corresponding author: Robbert Havekes Neurobiology expertise group, Groningen Institute for Evolutionary Life Sciences, University of Groningen, Groningen 9747 AG, The Netherlands.,: **Correspondence and requests** for materials should be addressed to R.H. equal contribution (Youri G. Bolsius, Pim R.A. Heckman). Department of Molecular, Cellular, and Developmental Biology, Ann Arbor MI 48109. **Author contributions** Author contributions R.H. conceived the study. Y.G.B., P.R.A.H., M.J.H.K., P.M., S.R., and R.H. designed the experiments. Y.G.B. and P.R.A.H. conducted the experiments. Y.G.B. and P.R.A.H. analysed the data. F.R. and E.L.M. assisted with data collection. Y.G.B., P.R.A.H., M.J.H.K., P.M., S.R., and R.H. interpreted the results. R.H., Y.G.B., and P.R.A.H. wrote the manuscript with input from M.J.H.K., P.M., S.R. M.J.H.K. P.M. and R.H. supervised the work.

## Abstract

It is well established that sleep deprivation after learning impairs hippocampal memory processes and causes amnesia. It is unknown, however, whether it leads to the actual loss of information or merely suppresses the retrievability of this information stored under suboptimal conditions. Here, we reveal that hippocampal memories formed under sleep deprivation conditions can be successfully retrieved multiple days following training using optogenetic memory engram activation or treatment with the clinically-approved phosphodiesterase 4 (PDE4) inhibitor roflumilast. Moreover, when optogenetic memory engram activation and roflumilast treatment were combined two days following training and subsequent sleep deprivation, it resulted in a more persistent memory trace that allowed for natural (*i.e.*, manipulation free) retrieval several days later. Our studies in mice demonstrate that sleep deprivation does not necessarily cause memory loss, but instead leads to the suboptimal storage of information that is difficult to retrieve. We also provide proof of principle that these suboptimally stored memories can be made accessible again far beyond the learning episode and that the clinically-approved PDE4 inhibitor roflumilast may be used to successfully retrieve information thought to be lost.

## Main document

Sleep loss is a common hallmark of modern society that affects people of all ages^1,2^ and has a severe impact on body and brain (*e.g.,*^3–9^). Even a single brief period of sleep deprivation following training can give rise to cognitive deficits, particularly in case of hippocampus-dependent memories^3,10–12^. Several misregulated signalling mechanisms have been identified that may contribute to these cognitive impairments^5,9–11,13–15^. Importantly, a crucial question that has not been addressed is whether sleep loss causes amnesia by preventing the storage of information, or by merely affecting the retrievability of information stored under suboptimal conditions. The development of novel approaches to label and activate neural ensembles forming a memory engram has greatly advanced our understanding of the molecular basis of memory, and allowed researchers to discriminate between cognitive problems arising from errors in encoding/consolidation (*i.e.,* information storage) and impairments in retrieval (*i.e.*, remembering) (e.g.,^16–18^). These findings have begun to reshape the ideas about how brain disorders as well as societal challenges affect brain plasticity critical for cognitive processes. In fact, these new insights led us to hypothesize that sleep deprivation does not lead to memory loss, but, instead, disrupts memory retrievability. To test this hypothesis, we applied optogenetic approaches and pharmacological strategies aimed at modulating memory engrams to restore access to information that might be hidden in the hippocampus.

## Sleep deprivation prevents the retrievability of newly formed memories

We examined whether sleep deprivation following learning attenuates the storage of information or merely hampers memory retrievability in adult male mice exposed to an object-location memory (OLM) task. This behavioural paradigm is based on the innate preference for spatial novelty (*i.e.*, the spatial relocation of an object), relies on the hippocampus^19^, and is susceptible to sleep deprivation^20–25^. Engram cells were successfully and specifically labelled when animals were taken off doxycycline (dox)-containing food from 24 hrs before OLM training until immediately after training (which corresponds with the start of the 6 hrs sleep deprivation period) (Fig. 1a-c). Labelling did not occur when animals were kept on dox (Fig. 1c). We performed a control experiment to examine whether mice instrumented for optogenetic manipulation would still have the expected preference for relocated objects in the OLM. Mice were maintained on dox throughout the experiment to prevent engram labelling, but were exposed to laser stimulation directly preceding the retrieval test (15 ms pulses of 20 Hz (473 nm) for 3 min at the level of the dentate gyrus (DG)) (Fig. 1d). The non-sleep deprived animals indeed showed a strong preference for the relocated object (Fig. 1e). In line with our previous work^20–25^, mice subjected to 6 hrs of sleep deprivation directly following training explored both objects to a similar extent indicating that they failed to detect the spatial novelty (*i.e.*, relocation of one object) (Fig. 1e; exploration times, Extended data Fig. 1a, b). We subsequently replicated the experiment but now took animals off dox 24 hrs preceding the training to allow for engram labelling (Fig. 1f). Optogenetic reactivation of this engram directly preceding the retention test successfully prevented the sleep deprivation-induced failure to detect spatial novelty (Fig. 1g; exploration times, Extended data Fig. 1c, d). Together, these findings suggest that sleep deprivation does not prevent the storage of information but, rather, suppresses the subsequent retrievability (*i.e.,* accessibility).

**Fig 1.**
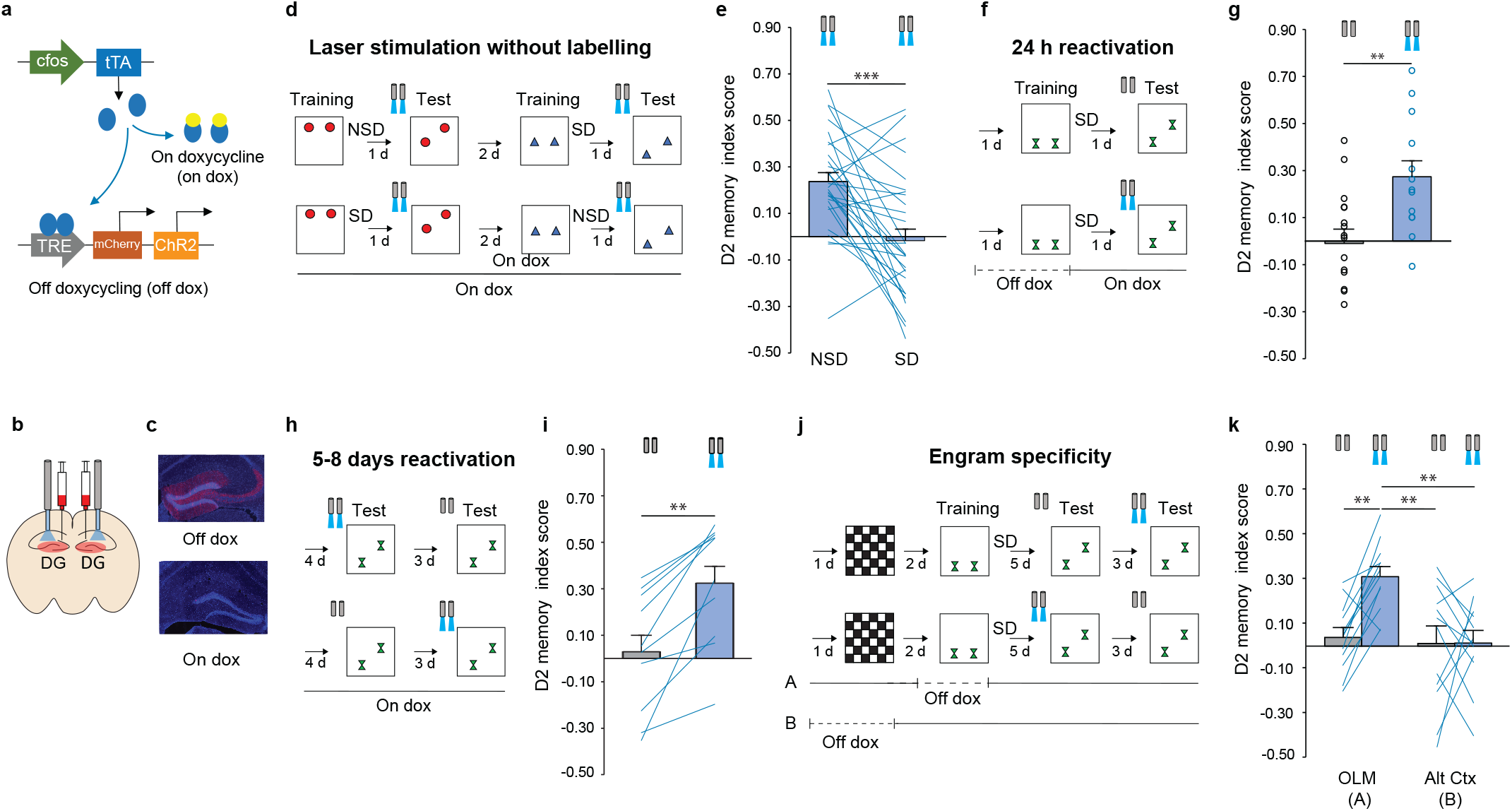
Sleep deprivation disrupts memory retrievability rather than causing the loss of information stored under sleep deprivation conditions. **a**, **b**, **c**, Mice expressing cfos-tTA were virally injected with AAV9-TRE-ChR2-mCherry and implanted with an optical fibre targeting the dentate gyrus (DG). Expression of FOS drives the synthesis of ChR2 & mCherry only in the absence of doxycycline (dox). **d**, Mice were trained in the object-location memory (OLM) task and subjected to 6 hrs of sleep deprivation directly following training or left undisturbed. All mice underwent laser treatment directly preceding the retention test. The experiment was replicated 2 days later with new objects and new object-locations in a cross-design fashion. Sleep deprivation group (SD), non-sleep deprived controls (NSD). **e**, Mice subjected to sleep deprivation failed to detect spatial novelty (*i.e.*, showed no preference for the relocated object) 24 hrs after training regardless of laser stimulation (both groups, *n*=29; paired samples t-test, *t*28=4.545; p<.001). **f**, Mice were taken off dox 1 day before OLM training to enable engram labelling, immediately put on back dox thereafter, subjected to 6 hrs of sleep deprivation, and tested 24 hrs after training. **g**, Engram reactivation, 24 hrs after training with sleep deprivation, resulted in the successful detection of spatial novelty (laser off, *n*=16; laser on, *n*=13; independent samples t-test, *t*27=-3.096; p=.005). **h,** Thereafter, mice received a second and third retention test at respectively 5 & 8 days after training with sleep deprivation. **i**, Engram reactivation 5 & 8 days after training with sleep deprivation led to the successful detection of spatial novelty (*n*=10; repeated measures ANOVA using ‘laser’ (on vs off) as within subject factor and ‘order’ (day 5 vs day 8) as between subject factor. No significant ‘laser*order’ effect was found (*F*1,8=.779; NS). Subsequent analysis showed a main effect for ‘laser’ (*F*1,8=25.250; p=.001). Additional paired samples t-test confirmed this effect of ‘laser on vs off’ (*t*9=-5.088; p=.001). **j**, To induce engram labelling, animals were taken of dox either during training in the OLM task or exposure to the alternative context (Alt Ctx). Engram reactivation was elicited 5 & 8 days thereafter. **k**, Reactivation of the OLM engram, but not Alt Ctx engram resulted in the successful detection of spatial novelty (all groups, *n*=12; repeated measures ANOVA using laser (on vs off) as within subject factor and context (OLM vs Alt Ctx) and order (laser ‘on-then-off’ vs ‘off-then-on’) as between subject factors. No three-way interaction (*F*1,20=.256; p=.618), or a laser*order interaction (*F*1,20=.000; p=.992) was found indicative that order of laser stimulation had no effect. A significant laser*context interaction was detected (*F*1,20=5.733; p=.027). Pairwise comparisons using Dunnett test with ‘OLM laser ON’ as reference against: 1) OLM laser OFF, p=.005; 2) Alt Ctx laser ON, p=.002; 3) Alt Ctx laser off, p=.002. All data are mean ± s.e.m. p*<.05, p**<.01, p***<.001.

Based on these observations, we wondered whether engram reactivation several days following training with sleep deprivation would still be able to restore memory retrieval and subsequent detection of spatial novelty in the OLM task. We therefore re-exposed the mice to two additional test sessions at 5 and 8 days following training with and without laser stimulation (Fig. 1h). Mice once again successfully detected the relocated object only if the retention test was directly preceded by laser stimulation (Fig. 1i; exploration times, Extended Data Fig. 1e). Laser stimulation on day 5 did not lead to subsequent novelty detection on day 8 (Fig. 1i). The latter observation is in line with previous work showing that optogenetic engram reactivation in itself is insufficient to permanently restore the retrievability of memory engrams^16,17,26^.

As a next step, we determined whether the successful retrieval was mediated specifically by reactivation of the OLM engram, or whether activation of any sparse population of neurons in the DG would lead to the same result. To label a subset of granular cells in the DG unrelated to the OLM task, we subjected a new batch of mice to an alternative context prior to training in the OLM task (Fig. 1j). Half of this group was taken off dox during exposure to the alternative context, whereas the other half was taken off dox during OLM training (Fig. 1j). Off dox exposure to the alternative context and OLM training resulted in a similar number of neurons labelled in the DG (Extended data Fig. 2). Following training and sleep deprivation, mice were subjected to the retention test 5 and 8 days thereafter with or without laser stimulation (Fig. 1k). Optogenetic activation of the neuronal population encoding the OLM resulted in a successful detection of spatial novelty (Fig. 1k; exploration times, Extended data Fig. 1f, g). In contrast, activation of the neural ensemble encoding the alternative context resulted in a failure to discriminate between the relocated and non-relocated object. These data indicate that activation of the neural ensemble encoding the OLM rather than a non-specific subpopulation of DG cells is required to overcome the sleep deprivation-induced failure to detect spatial novelty. Moreover, these findings highlight that sleep deprivation-induced impairments are a direct result of impaired memory retrievability of information stored under sleep deprivation conditions rather than the loss of information.

## Treatment with a PDE4 inhibitor is sufficient to overcome retrievability deficits caused by sleep deprivation

To increase the translational potential of our findings, we turned towards a non-invasive approach to overcome memory retrievability deficits. Previous work indicated that boosting cAMP signalling either systemically or locally in hippocampal excitatory neurons during sleep deprivation was sufficient to make memory processes resilient to the negative impact of sleep loss^20,27,28^. Therefore, we determined whether treatment with the clinically-approved phosphodiesterase 4 (PDE4) inhibitor roflumilast, to increase cAMP levels directly preceding the retention test, would lead to proper retrieval of memories stored under sleep deprived conditions. Twenty-four hrs after OLM training and sleep deprivation, mice were injected intraperitoneally (IP) with roflumilast (0.03 mg.kg^−1^) or vehicle solution 30 min preceding the test. Roflumilast treatment was sufficient to overcome the sleep deprivation-induced memory retrievability impairment (Fig. 2b; exploration times, Extended data Fig. 3a, b). Subsequently, we showed in a new batch of mice that we could successfully overcome retrieval deficits with roflumilast even 5 to 8 days after training (Fig. 2c, d; exploration times, Extended data Fig. 3c, d). Together, these data show that the clinically-approved drug roflumilast can successfully be used to successfully retrieve information stored under sleep deprivation conditions.

**Figure 2.**
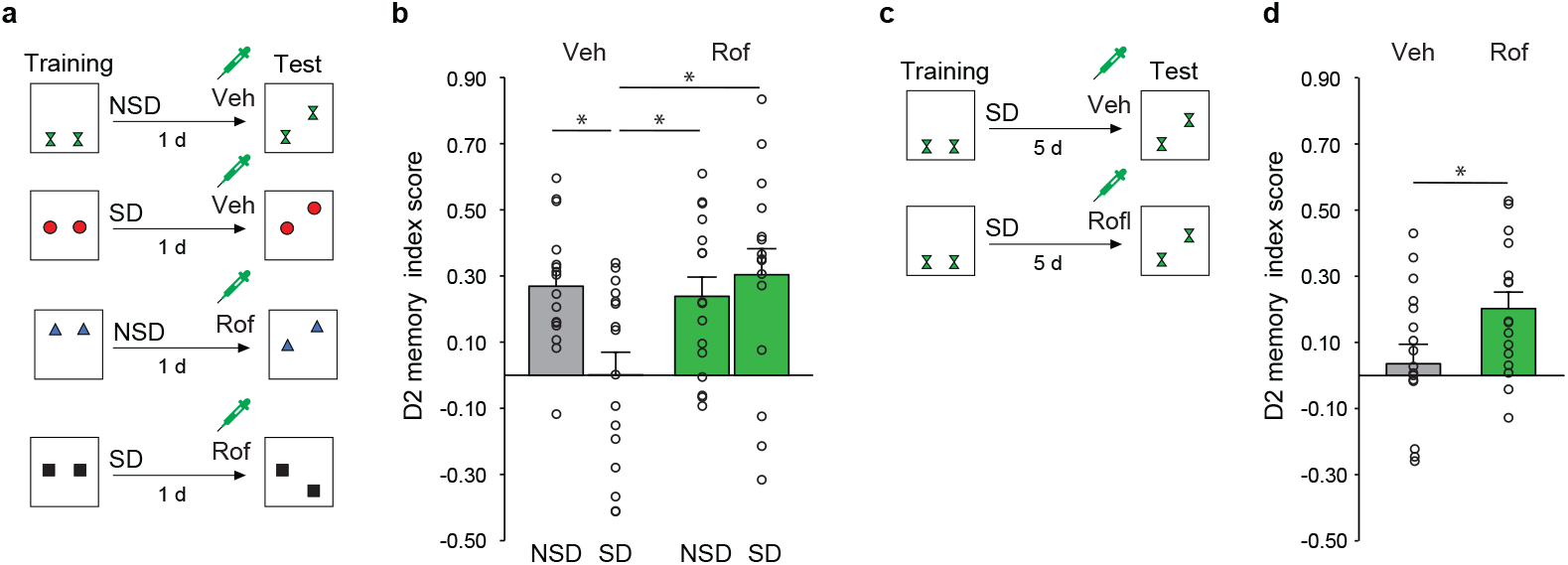
Treatment with roflumilast restores memory retrievability deficits caused by sleep deprivation following training. **a**, Following training in the OLM task, wild-type C57Bl/6 mice were subjected to 6 hrs of sleep deprivation (SD) or left undisturbed (NSD). Animals received an intraperitoneal (IP) injection with the PDE inhibitor roflumilast (rof; 0.03 mg.kg^−1^) or vehicle solution 30 min before the retention test 24 hrs after training. **b,** Roflumilast treatment preceding the retention test prevents memory retrieval deficits caused by sleep deprivation following training (all four groups, *n*=16, repeated measures ANOVA of one factor containing four levels, *F*3,45=3.928; p=.014, followed by pairwise comparisons using Dunnett post hoc test using ‘vehicle SD’ group as reference against which the other three groups are compared, ‘vehicle SD’ vs ‘Veh NSD’ (p=.012), vs ‘rof NSD’ (p=.03), vs ‘rof SD’ (p=.004). **c**, Following training in the OLM task, C57Bl/6 mice were subjected to 6 hrs of sleep deprivation. Animals received an IP injection with roflumilast (rof; 0.03 mg.kg^−1^) or vehicle solution 30 min before the retention test, 5 days after training. **d**, Roflumilast treatment preceding the retention test resulted in a proper detection of spatial novelty, 5 days after the training followed by sleep deprivation (both groups, *n*=16; independent samples t-test, *t*30=2.206; p=.035) All data are mean ± s.e.m. p*<.05, p**<.01.

## Restoring persistent access to previously irretrievable memories formed under sleep deprivation conditions

Optogenetic activation of hippocampal memory engrams only provides temporary access to information stored under suboptimal conditions (*e.g.*,^17,18^). Because transient short-term memories in rodents can be converted into more robust memories through systemic treatment with PDE4 inhibitors 3-5 hrs following training^29,30^, we combined optogenetic stimulation of the OLM engram with application of the PDE4 inhibitor roflumilast to test if we could make object-location memories stored under sleep deprivation conditions more persistently accessible and naturally retrievable. We trained a new batch of mice in the OLM task (and labelled neurons activated by learning as described (Fig. 1a-c)) and sleep deprived them for 6 hrs (Fig. 3a). Three days later, animals were re-exposed to the empty arena for 3 min and subjected to: 1) laser stimulation to activate the engram, 2) roflumilast treatment (0.03 mg.kg^−1^, IP; 3 hrs following training), or 3) a combination of laser stimulation and roflumilast treatment. Mice subjected to engram activation or drug treatment alone failed to detect the spatial novelty during the retrieval test 2 days later, *i.e.*, 5 days after the original training (Fig. 3a, b; exploration times, Extended data Fig. 4a, b). However, when optogenetic engram reactivation was combined with roflumilast treatment, animals were able successfully detect the spatial novelty two days after treatment (Fig. 3b; exploration times, Extended data Fig. 4a, b). Thus, while either optogenetic engram stimulation and drug treatment could provide short-lasting access to the information stored during sleep deprivation (*i.e.,* Fig. 1, 2), the combination of the two treatments resulted in a more permanent restoration of memory access that would allow for natural retrieval days after treatment without the need for any sort of stimulation or manipulation.

**Figure 3.**
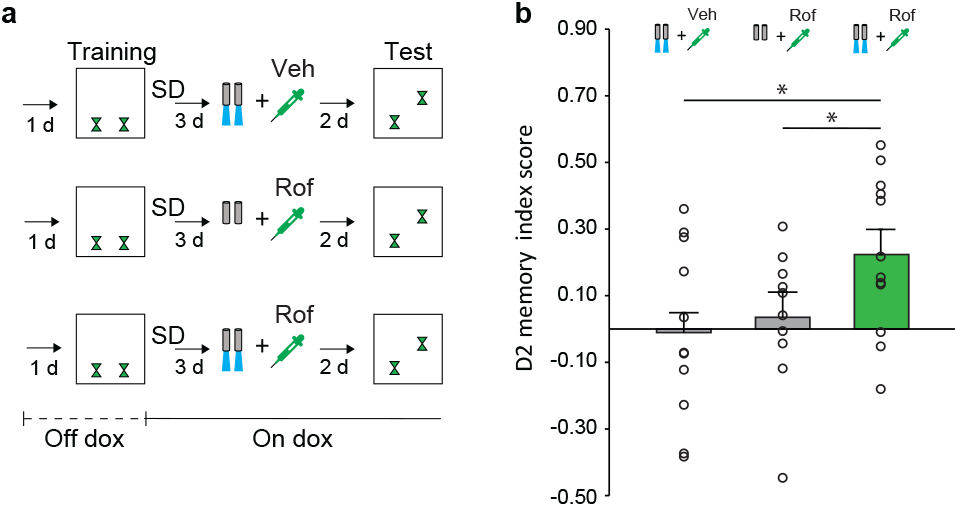
Restoring natural access to memories consolidated under sleep deprivation conditions. **a,** Mice expressing cfos-tTA were virally injected with AAV9-TRE-ChR2-mCherry and implanted with an optical fibre targeting the dentate gyrus. Mice were taken off dox 1 day before OLM training to enable engram labelling and immediately put on dox after training and subjected to 6 hrs of sleep deprivation (SD). Three days thereafter, mice were subjected to 1) optogenetic engram reactivation, 2) treatment with roflumilast (rof; 3 hrs after training), or 3) optogenetic engram reactivation in combination with roflumilast treatment. Mice were subjected to the retention test 2 days later (corresponding to 5 days after the training with sleep deprivation). **b**, Combining engram reactivation with roflumilast treatment resulted in the successful detection of spatial novelty, indicative of restored access to object-location memories consolidated under sleep deprivation conditions. Mice subjected to either engram reactivation or drug treatment failed to detect the spatial novelty (‘laser off+rof’, n=11; ‘laser on+veh’, *n*=10; ‘laser on+rof’, *n*=12; one-way ANOVA, *F*2,30=4.748; p=.016; Dunnett post hoc, ‘laser on+rof’ vs ‘laser off+rof’ (p=.017), and ‘laser on+veh’ (p=.032). All data are mean ± s.e.m. p*<.05, p**<.01.

In conclusion, our study reveals that deficits in hippocampal object-location memories consolidated under sleep deprivation conditions are not a result of the loss of information, but rather a disruption of the accessibility. Specifically, we show that object-location memories consolidated under sleep deprivation conditions can be successfully retrieved by optogenetic stimulation of the OLM engram or treatment with the PDE4 inhibitor roflumilast directly preceding the retention test. Intriguingly, these memories could be retrieved even up to 5 to 8 days after the learning event and sleep deprivation episode occurred. Most importantly, application of optogenetic engram stimulation in combination with roflumilast treatment days after training resulted in a more persistent memory trace that allowed for natural (*i.e.*, manipulation free) retrieval in the days thereafter.

While experiments have successfully retrieved memories under drug-based and pathologically-related conditions^17,18^, our study provides proof of principle data demonstrating that memories can be successfully retrieved and restored from sleep deprivation-induced amnesia. Indeed, a burgeoning literature suggests that memories thought to be “lost” still exist in a dormant state that can be artificially retrieved and behaviourally expressed^31,32^. One possibility is that the amnestic state renders a memory irretrievable by natural cues but in a manner that exogenous perturbations (e.g., optogenetics, drug treatment) can bypass, thereby providing evidence of successful memory retrieval. The conditions and molecular mechanisms underlying the retrievability of memories, despite the presence of amnesia, provide novel frameworks for studying the enduring changes experience imparts on the brain. Moreover, while previous studies successfully reactivated cells sufficient to produce freezing responses^33^, our study probes spatially modulated memories (*e.g.,* novel object locations) rather than freezing *per se*, which we believe opens up intriguing lines of research aiming to perturb such complex mnemonic processing, with a potential therapeutic role for the PDE4 inhibitor roflumilast.

## Methods

### Subjects

For the engram tagging studies (Fig. 1 and Fig. 3), we used hemizygous *c-fos-tTA/c-fos-shEGFP* male mice on a C57BL/6 background, which were bred in our facility (first breeding pairs from Jackson Labs). These transgenic mice have a tetracycline-transcriptional-transactivator (tTA) under the control of a c-fos promoter, originally developed by Dr. Mark Mayford. Pharmacological studies without light stimulation (Fig. 2) were done with C57BL/6 (Charles River labs). All mice were maintained on a 12-hrs light/12-hrs dark cycle and had *ad libitum* access to food and water. C*-fos-tTA/c-fos-shEGFP* were group housed prior to surgery, and were individually housed afterwards. Non-surgery C57BL/6 mice were group housed until one week prior to behavioural testing when they were individually housed. All mice were 2-5 months old during testing, which always took place at the beginning of the light phase. All mice initially received regular mouse chow (Altromin). However, one week prior to surgery, the c*-fos-tTA/c-fos-shEGFP mice* were put on food containing 40 mg.kg^−1^ doxycycline (dox) (Envigo). They remained on the dox diet until one day before the occurrence of the engram labelling. Mice were put back on regular chow 24 hrs prior to training in the learning task or exposure to the alternative context. Subsequently, the food was changed to a high dose dox chow (1 gkg^−1^) to prevent any further labelling and the animals were maintained on this diet throughout the experiment. All procedures were approved by the national Central Authority for Scientific Procedures on Animals (CCD) and the Institutional Animal Welfare Body (IvD, University of Groningen, The Netherlands), and conform Directive 2010/63/EU.

### Virus construct and packaging

In order to label memory engrams, we bilaterally injected an adeno-associated virus (AVV_9_-TRE-ChR2-mCherry, titre: 3.75e^14^, 200 nL; Penn Vector Core, Philadelphia, US) in the dentate gyrus (DG) of *c-fos-tTA/c-fos-shEGFP* mice using the following coordinates AP = 0.2, ML ± 0.13, DV = 0.18. Channelrhodopsin-2 (ChR2) and mCherry expression was controlled by the tetracycline response element (TRE). In this way, FOS expression would drive expression of tTA leading to the transcription of ChR2 and mCherry. The transcription of ChR2 and mCherry could be suppressed by keeping animals on a dox-containing chow (*i.e.*, c-fos expression would only lead to ChR2 and mCherry expression in the absence of dox).

### Viral injection and fibre optic implants surgery

All surgeries were conducted using a stereotaxic apparatus (Kopf instruments). Mice were anaesthetized with isoflurane (1.8–2.0%) and remained on a heating pad throughout the surgery. At the start of the surgery, mice were subcutaneously injected with carprofen (0.1 mL/10g), and 0.1 mL lidocaine was applied locally (subcutaneously) at the surgery site on top of the skull. Small bilateral holes were drilled in the skull using a 0.5 mm diameter microdrill at the appropriate locations. A 10 μL Hamilton micro syringe attached to a nanofil 33G needle (WPI) was loaded with virus and slowly lowered to the pocket site where it remained for one minute (see below for all coordinates). The needle was raised to the injection site where it again remained for one min before injecting started using a flow rate of 70 nL.min^−1^ and controlled by a microinjection pump (Micro4; WPI). A total of 200 nL virus was injected. After the viral injection, the needle remained at the target site for two min before it was slowly retracted from the brain. Optical fibres were mounted (see coordinates below), and stabilized with cement (C&B metabond) and two screws (0.96 mm diameter). At the end of the surgery, mice were injected with saline 0.1 mL (0.9%). All animals were allowed to recover for at least one week before starting behavioural experiments. The coordinates for the DG injections and optic fibre mounting were as follows, anterior-posterior (AP) −0.2 mm, medial-lateral (ML) ± 0.13 mm, dorsal-ventral (DV) −0.18 mm, and a pocket was made at DV −0.19 mm. The optic fibre was mounted at −0.2 mm AP and ± 0.13 mm ML, and the glass fibre was −0.175 mm long.

### Optogenetic stimulation protocol

Engram cells were optogenetically stimulated with 15 ms pulses of 20 Hz (473 nm) in accordance with previous work (pmid23888038). Stimulation was induced by means of a laser (CrystaLaser) which was driven by a TTL input (Doric lenses). The power delivered at the DG was 15-10 mW. All optogenetic laser stimulation was conducted in an open rectangular arena (empty OLM test arena) of 40 cm x 30 cm x 50 cm (length × width × height).

### Immunohistochemistry

Animals were transcardially perfused with 0.9% NaCl + Heparin and 4% paraformaldehyde in PBS (0.01M) followed by a 48hr post fixation at 4°C. Coronal brain sections were cut at a thickness of 20 microns. All sections were rinsed twice with PBS and incubated with 5% normal goat serum (NGS) in PBST (0.2% TritonX100) for one hour, followed by a 24 hrs incubation with primary antibodies at 4°C. After antibody incubation, all the sections were washed three times for 10 min with PBST and incubated for two hrs with the secondary antibodies diluted in NGS. A final incubation of 3 min with DAPI (1:1000, Thermo Fisher Scientific) was done, followed by three washing steps with PBST. Slices were mounted on glass slides and covered with mowiol (Sigma) to be visualized under a fluorescent microscope (Leica DMI6000). The following antibodies or combination of antibodies were used, mCherry [rabbit anti-mCherry, 1:1000, Invitrogen, LOT: TB254421; goat anti-rabbit Alexa 555, Thermo-fisher, LOT: A32730].

### Cell counting

For the quantification of the ChR2-mCherry expression in the DG, the number of positive mCherry cells were counted from 3-4 coronal hippocampal slices per animal. The coronal slices were taken from the dorsal hippocampus surrounding the coordinates covered by the optical fibre implants (−2.0 mm AP). Fluorescent images were acquired on a Leica DMI6000 using 10x magnification. Manual cell counting analysis was performed using ImageJ software. The cell body layer of dentate granule cells was marked as a region of interest based on the DAPI signal in each slice. For each section, the area was measured and the number of mCherry-positive cells was counted after subtraction of background fluorescence. Counting was performed blind to experimental condition.

### Sleep deprivation

Mice were sleep deprived during the first six hrs of the light phase, directly following the training trial of the object-location memory (OLM) task, using the gentle stimulation method (e.g.,^21,24,27^). In detail, animals were kept awake by gently tapping or shaking the cage. Shaking only occurred once tapping the cage was no longer sufficient. This sleep deprivation method has been extensively validated both at the behavioural level as well as through EEG recordings (*e.g.,*^34^), and work by us and others showed that behavioural and plasticity phenotypes associated with sleep deprivation were not caused by elevated plasma corticosterone levels or the gentle stimulation method itself (*e.g*.,^28,35–38^). More recently, we reported that blocking the synthesis and release of corticosterone in mice selectively during the SD period does not prevent sleep deprivation-induced memory deficits in the OLM task^24^.

### Drug preparation

The phosphodiesterase type 4 (PDE4) inhibitor roflumilast (Sigma Aldrich, Zwijndrecht, the Netherlands) was dissolved in vehicle solution containing 98% methyl cellulose Tylose (Sigma Aldrich, Zwijndrecht, the Netherlands) and 2% Tween80 (Sigma Aldrich, Zwijndrecht, the Netherlands) on the day of the behavioural studies, and injected in a volume of 2 mL.kg^−1^. Roflumilast was administered intraperitoneally at a dose of 0.03 mg.kg^−1^. Dose, injection volume, and injection schemes are based on our experience with the current drug in the OLM paradigm (*e.g.*, ^27,39^).

### Behavioural studies

#### Handling and habituation

Before the behavioural experiments started, all mice were handled and habituated for three consecutive days. On the first day, they were handled for 4 min/per mouse; on the second day mice were again handled 4 min/per mouse, and were habituated to scruff handling; on the third day, mice were brought to the experimental room and allowed to explore the empty arena for five min. Mice involved in the optogenetic experiments (Fig. 1 and Fig 3.) were connected to the optic fibre cable and allowed to explore the empty arena for five min with the laser turned on (15-10 mW, 15 ms pulses of 20 Hz).

#### Object-location memory (OLM) paradigm

This spatial learning task requires the hippocampus and is based on the innate preference for spatial novelty^19^. The OLM task was conducted in a rectangular arena (40 cm x 30 cm x 50 cm) with two spatial cues at the short walls on opposite sides of the arena (see ^22^ for a more detailed description of the arena and objects). The OLM task consisted of 10 min training and testing trials, during which the animals were allowed to freely explore two identical objects. During the training trial, the objects were placed symmetrically on a horizontal line, approximately 7.5 cm from the wall. In the testing trial, one of the objects was displaced along a straight line to a position that was 15 cm away from the training trial location. The combination of the side (left or right) and direction (up or down) of displacement of the objects in the testing trial was randomized and, during repeated testing, counterbalanced to avoid any place preferences. Object preferences were prevented by using different sets of objects for each animal in every experiment. Between animals and trials, the objects and arena were cleaned with a 70% ethanol solution to avoid the presence of olfactory cues. The exploration times per mouse for each object during the learning and test trial were manually scored using custom software (ORT v2; Maastricht, The Netherlands) by an experimenter blind to treatment. Directing the nose to the object at a distance of no more than 1 cm and/or touching the object with the nose was considered exploratory behaviour. From these exploration times, the relative measure of discrimination was calculated controlling for total exploration time: the *d2 index*. This relative discrimination index is calculated from the raw object exploration times during the test trial using the following equation: (exploration time object 1 at novel location - exploration time object 2 at original location) / (exploration time object 1 at novel location + exploration time object 2 at original location). For detailed description of the experimental design of each individual study, see supplementary information.

### Statistical analyses

IBM SPSS Statistics 26 was used to analyse all data. All statistical tests were two-tailed. All data was checked for normality. In addition, where applicable, Levene’s tests were first run to check for equality of variances. None of the conducted Levene’s tests were significant, therefore equal variances were always assumed. For all repeated measures ANOVA, the sphericity assumption was always checked using Mauchly’s sphericity test. Sphericity assumption turned out not to be violated in all analyses, meaning sphericity was assumed when inspecting tables showing tests of within-subject effects.

Data from Fig. 1g, 2d as well Extended data Fig. 1c, d, f, 2, and 3c, d was analysed using an independent-samples t-test. Data from Fig. 1e and Extended data Fig. 1a, b, were analysed using paired-samples t-tests.

Data from Fig. 1i was analysed by means of repeated measures ANOVA using ‘laser’ (on vs off) as within subject factor and ‘order’ (day 5 vs day 8) as between subject factor. No significant ‘laser*order’ interaction effect was found. Subsequent analysis of main effects showed a main effect for ‘laser’. An additional paired samples t-test confirmed this effect of ‘laser on vs off’.

For Fig. 1k, a repeated measures ANOVA was conducted using ‘laser’ (on vs off) as within subject factor and ‘context’ (OLM vs Alternative context (Alt Ctx)) as well as ‘order’ (laser ‘on-then-off’ vs ‘off-then-on’) as between subject factors. No three-way interaction was revealed, nor a laser*order interaction, both indicating that the order of laser stimulation had no effect and that the data of day 5 and 8 could validly be pooled. Finally, a significant laser*context interaction was shown. Next, we conducted pairwise comparisons using the Dunnett test with ‘OLM-context laser ON’ as reference against which all other conditions are compared: 1) against ‘OLM laser OFF’, 2) against ‘Alt Ctx laser ON’, and 3) against ‘Alt Ctx laser off’.

Data of Fig. 2b and Extended data Fig. 3a, b was analysed by means of repeated measures ANOVA using one within factor containing four levels (*i.e.*, the four experimental conditions). After showing significance of the overall ANOVA for Fig 2b, we continued with a pairwise comparisons by means of Dunnett post hoc test using ‘vehicle SD’ group as reference against which the other three groups are compared, ‘veh SD’ vs ‘Veh NSD’, vs ‘rof NSD’, vs ‘rof SD’. For Extended data Fig. 3a, b no significant interaction effect was observed.

For Fig. 3 and Extended data Fig. 4a, b, a one-way ANOVA was used with ‘experimental condition’ as between-subject factor containing three levels. After showing significance of the overall ANOVA for the data in Fig. 3, additional post hoc analyses using Dunnett showed that both ‘laser off+rof’, and ‘laser on+veh’ differed from the reference group ‘laser on+rof’. For the Extended data Fig. 4a, b no significant interaction effect was observed.

Extended data Fig. 1g was analysed using repeated measures ANOVA using ‘laser’ (on vs off) as within subject factor and ‘context’ (OLM vs Alt Ctx) as between subject factor. No significant interaction or main effect was observed.

## Data availability

All data are available from corresponding authors upon reasonable request.

### Acknowledgements

We would like to thank members of the Havekes lab for help with the sleep deprivation studies and Dr. Tomas Ryan for fruitful discussions at the start of the project. This work was supported by the Human Frontiers Science Program Organization (HFSP) (Grant RGY0063/2017 to RH), and Netherlands Organisation for Scientific Research (Grant NWA.1228.191.208 to R.H. and S.R.).

## Notes

**Competing interests** The authors declare no competing interests

### Competing Interest Statement

The authors have declared no competing interest.

