## supplementary methods and extended data legends for "Restoring persistent accessibility to memories after sleep deprivation-induced amnesia"

**Supplementary Information**

Extended description of all behavioural training and testing protocols.

*Sleep deprivation and laser stimulation and the object-location memory task* (Fig. 1d, e; Extended data Fig. 1a, b)*.* In this experiment, we examined whether sleep deprivation following training attenuated memory performance during the test trial 24 hrs after learning. Moreover, we investigated whether optogenetic stimulation hampers the normal learning and formation of object-location memories. Mice were kept on a dox diet (40 mgkg^-1^) throughout the whole experiment to prevent any tagging. First, mice were handled and habituated according to our handling and habituation protocol (see the methods section). They underwent an object-location memory (OLM) training trial, and were immediately sleep deprived (SD) or left undisturbed (NSD) depending on their experimental condition (Fig. 1d). The testing procedure 24 hrs later was according to the optogenetic testing trial protocol (Extended data Fig. 5). *The optogenetic testing trial protocol* was as follows*,* mice were first connected to the fibre optic cable and placed back into their home cage for 90 sec Next, they were placed into the empty arena *without objects* for three minutes. During these three min, animals received laser stimulation. Subsequently, after the laser stimulation, the mice were placed back into their home cage for five min, before being placed into the arena *with objects* for ten min (testing trial). Two days after the testing procedure, we repeated the whole experiment including training and test trial. However, during this second experiment, we switched experimental groups in such a way that animals that were sleep deprived during the previous session were now left undisturbed, and *vice versa*. Data of both experiments was pooled so that every mouse constitutes its own control (*i.e.*, sleep deprived versus normal sleep).

*Engram reactivation 24 hours after learning episode followed by sleep deprivation* (Fig. 1f, g; Extended data Fig. 1c, d*).* For this experiment, we used the same mice as in the *optogenetic stimulation without engram labelling* study, described above. Importantly, different objects and locations were used to avoid any carry-over effects. In this experiment, the engram of the training trial was tagged by replacing the dox diet (40 mg.kg^-1^) with normal chow 24 hrs before the training trial. To prevent any tagging after the training trial, we exchanged the normal chow with a high-dose dox diet (1g kg^-1^) immediately after the training trial.

During the training trial, animals were allowed to explore two similar objects for ten min. Immediately following training, all animals were sleep deprived for six hrs. Twenty-four hours after the training trial, animals underwent a test trial according to our *optogenetic testing trial protocol* (Extended data Fig. 5). Mice were first connected to the laser and placed back into their home cage for 90 sec. Next, they were placed into the empty arena *without objects* for three minutes. During these three min, half the animals received laser stimulation, while the other half received no laser stimulation. Subsequently, after the laser stimulation, the mice were placed back into their home cage for five min, before being placed into the arena *with objects* for ten min (testing trial).

*Engram reactivation 5 & 8 days after learning episode followed by sleep deprivation*. (Fig. 1h, I; Extended data Fig. 1e). In order to investigate whether the memory engram could still be optogenetically reactivated after a prolonged period, we used the same mice as in the 24 hrs reactivation experiment described above. Upon completion of the testing trial of the 24 hrs reactivation experiment, animals were left undisturbed for four days. Thus, five days after the training trial followed by sleep deprivation, mice underwent an additional delayed testing trial. The procedures for the current testing trial were similar to those described for the testing trial of the 24 hrs reactivation experiment (Extended data Fig. 5) with the important distinction that the group receiving laser stimulation now received no laser stimulation, and vice versa. Three days after this delayed reactivation trial, we subjected the mice to another testing trial, during which we again interchanged laser stimulation conditions.

*Reactivation OLM engram versus alternative context engram* (Fig. 1j, k; Extended data Fig. 1f, g). This experiment was conducted to demonstrate that our behavioural data was induced specifically due to engram reactivation of the tagged learning episode, and is not induced by activation of an engram from an unrelated learning episode. Mice were first habituated according to our habituation protocol (see the methods section). After habituation they were divided into two groups: training context versus alternative context. Both groups were subjected to similar experimental procedures, and only differed in their "tagged context", which was accomplished by taking the mice off dox food at different timepoints during exposure to different contexts. Both groups were first exposed to an alternative context (day 1), followed by the exposure of an empty arena 24 hrs later (day 2). On day 3, they underwent the OLM training trial. Whereas one group was taken off dox 24 hrs before the OLM training trial, the other was taken off dox 24 hrs before exposure to the alternative context (Fig. 1j). Both groups were deprived of sleep after the OLM training trial, and transferred back to the housing room thereafter where they were left undisturbed for the next 5 days. On day 5, the tagged engram was reactivated and tested according to the optogenetic testing protocol (Extended data Fig. 5). All groups were divided in such a way that ‘reactivation’ or ‘no reactivation’ conditions were balanced in both the training context and alternative context group. Three days following the testing trial, all animals were subjected to another retention trial, however, the reactivation conditions were reversed between groups in a cross-design fashion.

*Treatment with roflumilast 24 hrs after OLM training followed by sleep deprivation* (Fig. 2a, b; Extended data Fig. 3a, b). In this study, no optogenetic stimulation was applied; only drug treatment with roflumilast. Mice were first habituated and handled according to our handling protocol (see the methods section), with the addition of an intraperitoneal (IP) injection with saline (0.03 mL per animal) during the last day of handling. Twenty-four hrs later, on experimental day 1, mice underwent a training trial in which they explored two similar objects for 10 min. The six hrs of sleep deprivation took place immediately after the training trial, while the non-sleep deprived animals were left undisturbed. The testing trial took place 24 hrs after the training trial, on day 2. All mice received a single injection with either roflumilast or vehicle 30 min before the testing trial.

*Treatment with roflumilast 5 days after OLM training followed by sleep deprivation* (Fig. 2c, d; Extended data Fig. 3c, d*).* For the current experiment, we first handled the animals according to our handling protocol (see the methods section) with the addition of an IP injection with saline (0.03 mL per animal) during the last day of handling. On experimental day 1, mice underwent a training trial in which they explored two similar objects for 10 min. Thereafter, all the groups were sleep deprived for 6 hrs. After 5 days, the testing trial took place. Mice were first exposed to the box *without objects* and placed back in the home cage. Next, they were subjected to a testing trial in which one of the objects was relocated to a novel location. The mice were injected with vehicle solution or roflumilast (depending on the experimental group) 30 min before the exposure to the empty box.

*Optogenetic engram reactivation in combination with roflumilast treatment to restore the accessibility of spatial memories under sleep deprived conditions* (Fig. 3; Extended data Fig. 4a, b). Similar to all previously described studies, mice were first handled and habituated (see the methods section). At the end of the third habituation day and 24 hrs before the training trial, mice were taken off dox food. On experimental day 1, mice underwent the learning trial in which they were allowed to explore two similar objects for 10 min. After the training trial, we immediately changed the diet to a 1 gkg^-1^ dox diet, and subjected them to 6 hrs of sleep deprivation. The next 3 days, the animals were left undisturbed. On day 4, the reactivation session took place, and the mice were divided into three experimental conditions: ‘laser on + vehicle’, ‘laser off + roflumilast’, or ‘laser on + roflumilast’. The reactivation session was in the OLM arena *without objects* and lasted three min. During these three min the laser was turned on in the laser on experimental groups. Three hrs after the reactivation we injected roflumilast or vehicle IP. Two days later mice were subjected to a “normal” testing session in which one of the objects was moved to a novel location. During this test session, no drugs or laser treatment was applied.

**Extended data figure legends**

**Extended data figure 1. Object exploration times for the training and testing trials of the engram reactivation studies.** **a**, total object exploration times of the training trial for the study in which all animals were kept on dox throughout the experiment but did receive laser stimulation (related to Fig. 1d). During this experiment mice were sleep deprived (SD) or not (NSD) directly following training. Twenty-four hrs after the training trial they were subjected to the testing session. The NSD and SD groups did not differ in their object exploration times during training (NSD, *n*=29; SD, *n*=29, paired samples t-test, *t*_28_=.158; p=.875). **b,** Similarly, total exploration times did not differ for the same experiment between NSD and SD conditions during the testing trials (NSD, *n*=29; SD, *n*=29, paired samples t-test, *t*_28_=.006; p=.995). **c,** The next experiment related to the reactivation of the memory engram 24 hrs after learning aimed at reversing the SD-induced memory deficit. All mice were sleep deprived immediately after the training trial. The laser on and laser off groups did not differ in their total exploration times of the objects during the training trials (laser off, *n*=16; laser on, *n*=13, independent samples t-test, *t*_27_=-.298; p=.768). **d**, Also during the subsequent testing session, laser on and off conditions did not alter the object exploration times (laser off, *n*=16; laser on, *n*=13, independent samples t-test, *t*_27_=-.668; p=.510). **e**, When the mice were subjected to an additional testing trial, 5 and 8 days after the initial learning experience, the laser on and laser off groups did not differ in their total object exploration times (laser off, *n*=10; laser on, *n*=10, paired samples t-test, *t*_9_=-.099; p=.924). **f**, The engram specificity study in which dentate gyrus (DG) neurons were labelled during the object-location (OLM) training or alternative context (Alt Ctx) exposure, and subsequently reactivated 5 and 8 days later. Importantly, again no differences in total object exploration times were observed between groups (OLM, *n*=24; Alt Ctx, *n*=24, independent samples t-test, *t*_22_=-1.201; p=.243). **g**, To analyse the exploration times of the test trials, we used a repeated measures ANOVA with laser (on vs off) as within subject factor and context (OLM vs Alt Ctx) as between subject factor. The ANOVA showed no significant interaction (all groups *n*=12, repeated measures ANOVA, *F*_1,22_=.011; p=.916). Subsequent inspection of main effects revealed no significant main effect for laser (*F*_1,22_=0.129; p=0.723). Of note, Levene’s test for equality of variances was not significant for all analyses. Therefore, where applicable, equal variances were always assumed. Taken together, these data indicate that our experimental findings cannot be explained by changes in the total object exploration times during the training or testing trials. Therefore, these data further support the notion that sleep deprivation impacts the retrievability of information when stored under sleep deprived conditions. All data are mean ± s.e.m.

**Extended data figure 2. Quantification of the ChR2-mCherry positive cells in the dentate gyrus neuronal ensemble of the object-location engram and alternative context engram.** In this study, the object-location memory (OLM) engram or the alternative context (Alt Ctx) engram were labelled and reactivated 5 and 8 days after the OLM training trial followed by sleep deprivation (SD). Two groups, the OLM and Alt Ctx groups, were both exposed to the Alt Ctx for a duration of 10 min and, 48 hrs later, to an object-location training trial, again for 10 min. However, in the OLM group, the neuronal ensemble in the dentate gyrus (DG) was labelled during the OLM training trial, while in the Alt Ctx group, the Alt Ctx was labelled. The amount of positive mCherry-ChR2 cells in the DG were counted (3-4 hippocampal slices per animal) in both groups to determine whether exposure to an alternative context and OLM training resulted in a similar number of tagged cells. The number of tagged engram cells did not differ between groups (OLM, *n*=8; Alt Ctx, *n*=9, independent samples t-test, *t*_15_=.108; p=.915). These data indicate that the successful memory retrieval after reactivation of the neural ensemble labelled following OLM training could not be explained by potential differences in the number of cells labelled in the DG during OLM training and alternative context exposure.

**Extended data figure 3. Object exploration times during the training and testing trials of the roflumilast studies.** **a**, Exploration times of the training trials belonging to the studies in which the sleep deprivation (SD)-induced retrievability deficit was rescued by treatment with the PDE4 inhibitor roflumilast (rof). In this experiment, mice were subjected to a training trial immediately followed by SD or normal sleep (NSD). None of the four experimental groups differed in their exploration time during the training trial (all four groups, *n*=16, repeated measures ANOVA of one factor containing four levels, *F*_3,45_=.797; p=.502). **b**, After 24 hrs, animals underwent the test trial. Thirty min prior to testing, the animals received either roflumilast (rof) or vehicle (veh). The object exploration times of the testing trials did not differ between the four experimental groups (all four groups, *n*=16, repeated measures ANOVA of one factor containing four levels, *F*_3,45_=1.123; p=.350).**c,** In the subsequent experiment, we tested in a separate batch of mice whether the retrievability of the memory that is lost due to SD could still be rescued by roflumilast when activated after a prolonged period of five days. To this end, all mice were sleep deprived after the training trial, and were treated with roflumilast or vehicle 30 min before testing. The vehicle and roflumilast groups did not differ in their object exploration times during training (both groups, *n*=16; independent samples t-test, *t*_30_=-.668; p=.510). **d**, Similar to the training trail, the vehicle and roflumilast groups did not have different object exploration times when they were tested 5 days later (both groups, *n*=16; independent samples t-test, *t*_30_=.712; p=.482).

**Extended data figure 4. Exploration times of the objects during the training and testing trials of the restored natural access studies.** In this experiment, we tagged the engram of the object-location memory (OLM) training trial in order to restore the natural access to memories stored under sleep deprived conditions. This was established by combining an engram reactivation session with treatment with the phosphodiesterase 4 (PDE4) inhibitor roflumilast (rof), 3 days after the OLM training trial followed by sleep deprivation (SD). **a**, The exploration times of the training trials 3 days before the reactivation session did not differ between groups receiving laser stimulation or roflumilast alone, or their combination (laser off + rof, *n*=11; laser on + veh, *n*=10; laser on + rof, *n*=12, one-way ANOVA, *F*_2,30_=.479; p=.624). **b**, Similarly, the exploration times during the testing trials did not differ between the three different conditions (laser off + rof, *n*=11; laser on + veh, *n*=10; laser on + rof, *n*=12, one-way ANOVA, *F*_2,30_=1.396; p=.263). Together, this data contributes to the evidence showing that the restored access to memories consolidated under sleep deprived conditions cannot be explained by any differences in total object exploration times during the training or testing trials.

**Supplementary figure 5. Overview optogenetic testing protocol.** In all optogenetic testing stimulation sessions we followed this protocol. Mice were first connected to the laser and placed in their homecage for 90 sec. Thereafter, they were placed in the experimental box (length 40 cm x width 30 cm x height 50 cm) without objects for 3 min. During these 3 min the laser manipulation took place (15 ms, 20 Hz, 10-15 mW). After laser stimulation, mice were placed back in their homecage for 5 min. Lastly, mice were subjected to the testing trial for 10 min in which they were placed again in the experimental box, but this time with objects. One of the objects was moved to a novel location compared to the initial training trial.
