## Supplementary figures and images for "Restoring persistent accessibility to memories after sleep deprivation-induced amnesia"

### Extended data figure 1

Bolsius, Heckman et al,  Extended data, Figure 1

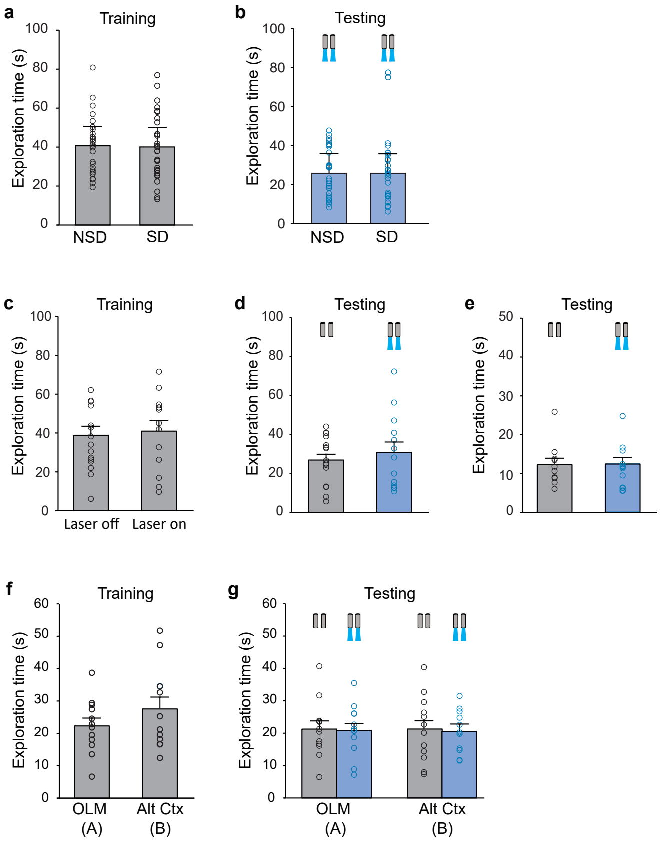

### extended data figure 2

Bolsius, Heckman et al,  Extended data, Figure 2

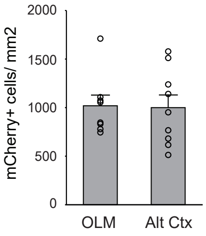

### extended data figure 3

Bolsius, Heckman et al, Extended data, Figure 3

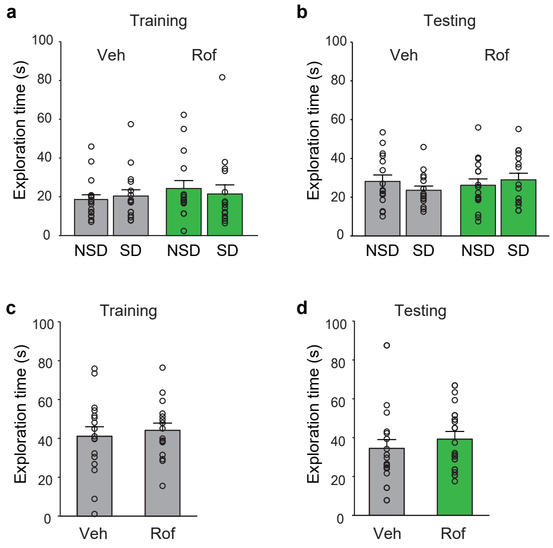

### extended data figure 4

#### Bolsius, Heckman et al,  Extended data, Figure 4

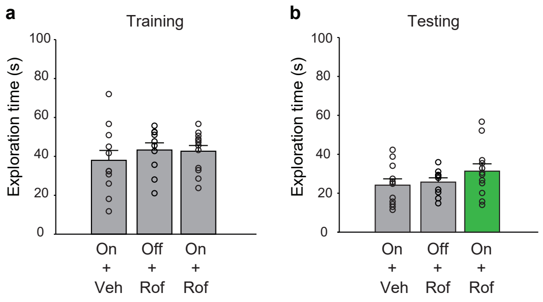

### extended data figure 5

Bolsius, Heckman et al,  Extended data, Figure 5

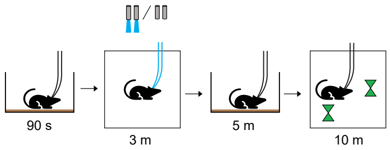
